# Comparison of Pairwise Samples of Endotracheal Aspirate and Bronchoalveolar Lavage for Microbial Analysis in Critically ill Patients

**DOI:** 10.1101/2025.10.08.680399

**Authors:** Sina Pesthy, Silvia Gschwendtner, Michael Schloter, Nadine Liebchen, Teresa Kauke, Julia Kovács, Uwe Liebchen

## Abstract

**Introduction:** Bronchoalveolar lavage (BAL) is considered as gold-standard for the characterization of the lung microbiome. The aim of this study was to elucidate the applicability of less-invasive sampling by endotracheal aspirate (ETA) compared with BAL for the analysis of respiratory microorganisms in critically ill patients who required mechanical ventilation.

**Methods:** Pairwise samples of ETA and BAL obtained from mechanically ventilated patients at our intensive care units were collected anonymously for testing the accuracy of ETA compared to BAL. Bacterial community structure was assessed by a metabarcoding approach based on 16S rRNA gene sequencing in ETA and BAL samples.

**Results:** In total, 13 samples of each, BAL and ETA, were collected from 13 critically ill patients. No differences between BAL and ETA were found for alpha diversity based on species richness (*p*=0.77), Shannon diversity (*p* = 0.41), Simpson index (*p* = 0.85) and evenness (*p* = 0.98). Overall, BAL and ETA samples demonstrated strong taxonomic concordance at the levels of bacterial phyla and genera but differed distinctly at the amplicon sequence variants (ASV) level.

**Conclusion:** In an unselected cohort of mechanically ventilated patients, BAL and ETA samples exhibited profound resemblance at higher taxonomic ranks, with increasing divergence observed at finer taxonomic resolutions. Our findings may facilitate guidance towards the reliability of non-invasive ETA as a valuable approach for clinical studies investigating the lung microbiome.

## INTRODUCTION

The lower respiratory tract has historically been acknowledged as a sterile compartment, primarily based on the absence of resident pulmonary microbiota in culture-based studies [1]. However, this concept of lung sterility was fundamentally challenged by the implementation of culture-independent technologies for microbial detection [2]. High-throughput DNA sequencing discovered a dynamic population of microbiota within the bronchial tree of the lung harboring approximately 2.2 × 10^3^ bacterial genomes per cm^2^ [3,4]. In fact, this microbial equilibrium of the lung is continuously exposed to environmental factors through gas exchange and microaspiration with gut-to-lung translocation of microbiota [5–8]. In healthy individuals, these exogenous and endogenous factors contribute to a dynamic and constantly adapting microbial ecosystem [7]. Conversely, disruption of this balance may lead to pulmonary dysbiosis with elevated bacterial burden potentially perpetuating alveolar inflammation [9].

The collection of lower respiratory secretion is substantial for the precise characterization of the lung microbiome and thereby elucidating the contribution of microbial communities to respiratory pathology, especially in mechanically ventilated critically ill patients [3,9– 11].

Among the most common sampling techniques for the procurement of specimens of the lower respiratory tract are bronchoalveolar lavage (BAL) and endotracheal aspirate (ETA). Acquisition of BAL is an invasive, time-consuming and cost-intensive technique demanding specialized expertise and equipment. In contrast, ETA is a non-invasive and easily accessible method in routine clinical practice, facilitating safe and convenient serial sampling in ventilated patients for longitudinal microbiome assessment.

The diagnostic value of ETA has traditionally been considered inferior to BAL since ETA-based culture analysis may be affected by a lack of specifity due to potential contamination with commensal microorganisms from the oropharynx and upper respiratory tract [12–15]. Consequently, bronchoscopic BAL remains the preferred approach for culture-based pathogen detection in clinical practice, despite its increased risk of procedure-related complications, e.g. hemorrhage, hypoxemia and sedation-associated risks, as well as logistical challenges, such as ensuring round-the-clock availability of bronchoscopes and skilled personnel in the intensive care setting [12,16]. In this context, the reliance on invasive sampling methods such as BAL for sequence-based analysis of the respiratory microbiome in intubated patients should be point of discussion.

Thus, the aim of this study was to elucidate the applicability of non-invasive ETA sampling compared with BAL for analysis of microorganisms in the lower respiratory tract of unselected critically ill patients based on gene sequencing techniques.

## METHODS

### Study design

In total, 13 pairwise ETA and BAL samples from 13 critically ill patients were included in this study. All samples were obtained anonymously at the intensive care unit of the LMU University Hospital Munich during routine clinical care or standard diagnostic evaluations without any additional procedures performed for research purposes. ETA and BAL samples were procured solely in accordance with clinical practice by trained and experienced intensive care physicians at our institution. The local ethic committee has granted approval for publication of this anonymous, retrospective study (Ethics Committee of the Medical Faculty of LMU Munich, Project No. *25-0228*, Date: *20*.*03*.*2025*).

BAL was performed under guidance of a flexible fiberoptic bronchoscope (BF-1 TH 190 or BF-1 TH 1100, Olympus Europa SE & Co.KG, Hamburg, Germany) or a single-use device (Ambu^®^ aScope^™^ 5 Broncho HD 5.0/2.2, Ambu A/S, Ballerup, Denmark), as clinically recommended. Following the passing of the catheter through the endotracheal or tracheostomy tube and advancing into the right or left main bronchus, saline lavage was conducted with a volume of 10mL, subsequently reaspirated and collected in a sterile tube. The collected fluid was resuspended carefully to achieve homogeneity preliminary to be frozen within one hour and stored at −80°C until further research. ETA was obtained via tracheal suctioning by utilizing a 14 Fr Closed Suction System (Ballard Closed Suction System, Avanos Medical Deutschland GmbH, Hamburg, Germany) homogenized and preserved at −80°C until microbiological analysis.

### Sequence library processing

1mL of BAL and ENTA samples were centrifuged at 20,000 g or 10 min at 21°C before extracting the DNA from pelleted cells via NucleoMag DNA Forensic kit (Macherey Nagel) according to manufacturers’ instruction. Amplicon sequencing of the V3-V4 hypervariable region of the 16S rRNA gene was performed on a MiSeq Illumina instrument (MiSeq Reagent Kit v3 (600 Cycle); Illumina, San Diego, CA, USA) using the universal eubacterial primers 347F and 803 [17], extended with sequencing adapters to match Illumina indexing primers. To identify potential contamination during DNA extraction and amplification, both extraction and PCR no template control samples were performed. PCR was done using NEBNext high fidelity polymerase (New England Biolabs, Ipswich, USA) in a total volume of 25 µl (10 ng DNA template, 12.5 µl polymerase, 5 pmol of each primer). PCR conditions were 5 min at 98 °C; 32 cycles of 10 seconds (sec) at 98 °C, 30 sec at 56 °C, 30 sec at 72 °C; 10 min 72 °C. PCR products were purified using MagSi NGSprep Plus beads (Steinbrenner, Wiesenbach, Germany) and quantified via PicoGreen assay. Subsequently, indexing PCR was performed using the Nextera XT Index Kit v2 (Illumina, Inc. San Diego, CA, US) in a total volume of 25 µl (10 ng DNA template, 12.5 µl NEBNext high fidelity polymerase, 2.5 µl of each indexing primer) and the following PCR conditions: 30 sec at 98 °C; 8 cycles of 10 sec at 98 °C, 30 sec at 55 °C, 30 sec at 72 °C; 5 min 72 °C. Indexing PCR products were purified using MagSi NGSprep Plus beads, qualified and quantified via a Fragment Analyzer™ instrument (Advanced Analytical Technologies, Inc., Ankeny, USA) and pooled in an equimolar ratio of 4nM.

FASTQ files were trimmed with a minimum read length of 50 using Cutadapt [18] followed by quality control via FastQC [19]. For subsequent data analysis, DADA2 version 1.26 [20] was used with the following trimming and filtering parameters: 20 bp were removed n-terminally, and reads were truncated at position 280 (forward) and 280 (reverse), respectively, with an expected error of 6 (forward) and 8 (reverse). With respect to rare variants the pseudo-pooled sample processing approach was chosen, which includes two rounds of independent sample processing, whereby the second round is trained by “prior” sequence variants from the first round (sequences which could be expected). This approach increases sensitivity to rare variants without an increase in spurious sequences. The resulting unique amplicon sequence variants (ASV) were assigned to the SILVA v138.1 (release 99%) reference database. Reads were excluded if classified as mitochondria or if the phylum was missing. In addition, singletons (ASV represented by only one read) were removed from the dataset. All blank extraction and PCR no template controls were analyzed together with biological samples, but no potential contaminants were identified by the ‘decontam’ R package [21]. Thus, all 588 ASV were used for subsequent analysis.

### Data analysis

All plots and statistics were performed in R version 4.3.2 (https://www.R-project.org). Sequencing data were normalized using the Trimmed Mean of M-values method [22]. Alpha diversity was calculated using species richness, evenness as well as Shannon and Simpson diversity index. Beta diversity was analyzed via Bray Curtis and Jaccard distance. For statistical purposes, generalized linear mixed-effects models including the study subjects as random factor by defining a random effect term (R package nlme and lme4) and PERMANOVA using strata to restrict permutations within blocks (R package vegan) were used. P value adjustment for multiple comparisons was performed with Benjamini-Hochberg correction. Plots were created using the packages phyloseq, ggplot2 and ggpubr. Shared and unique ASV were calculated using the packages microbiome.

Alpha diversity and Venn diagrams of ASV were visualized using GraphPad Prism (GraphPad Prism software, Version 9).

## RESULTS

In total, 13 samples of each, BAL and ETA, were obtained from 13 mechanically ventilated patients from the intensive care unit at LMU University Hospital Munich. Demographic and clinical data of enrolled patients was not assessed due to the anonymous character of this study.

A total of 830,994 raw reads were obtained in BAL and 832,921 unfiltered reads in ETA. Subsequent to quality filtering for unassigned reads, 801,120 reads remained for BAL and 802,524 reads for ETA. Adjustment of raw reads for human-associated sequences (e.g. mitochondrial) resulted in 423,574 filtered reads for BAL and 438,563 reads for ETA. Solely two singleton reads were detected in ETA-derived samples and excluded from further analysis, as appropriate. It should be noted that filtering for human sequences resulted in low percentage of present reads in *subject 1*, though rates were comparable between BAL (1.2%) and ETA (1.1%).

With respect to alpha diversity Shannon, Simpson and Evenness Index were 1.3 (*IQR* 0.7–2.6; *range*[*min*–*max*] 0.09–3.0), 0.5 (*IQR* 0.4–0.9; *range*[*min*–*max*] 0.03–0.9) and 0.5 (*IQR* 0.3–0.7; *range* [*min*–*max*] 0.03–0.8), respectively in BAL compared to 1.4 (*IQR* 0.7– 2.1; *range*[*min*–*max*] 0.2–2.9), 0.7 (*IQR* 0.4–0.8; *range*[*min*–*max*] 0.08–0.9) and 0.4 (*IQR* 0.3–0.6; *range*[*min*–*max*] 0.09–0.8) in ETA.

Collectively, no difference in Shannon diversity (*p* = 0.41), Simpson index (*p* = 0.85) and evenness index (*p* = 0.98) between the pairwise samples of BAL and ETA was observed, as demonstrated in **Figure 2B-D**. Thus, these results indicate a similar alpha diversity in ETA and BAL samples.

**Figure 1.**
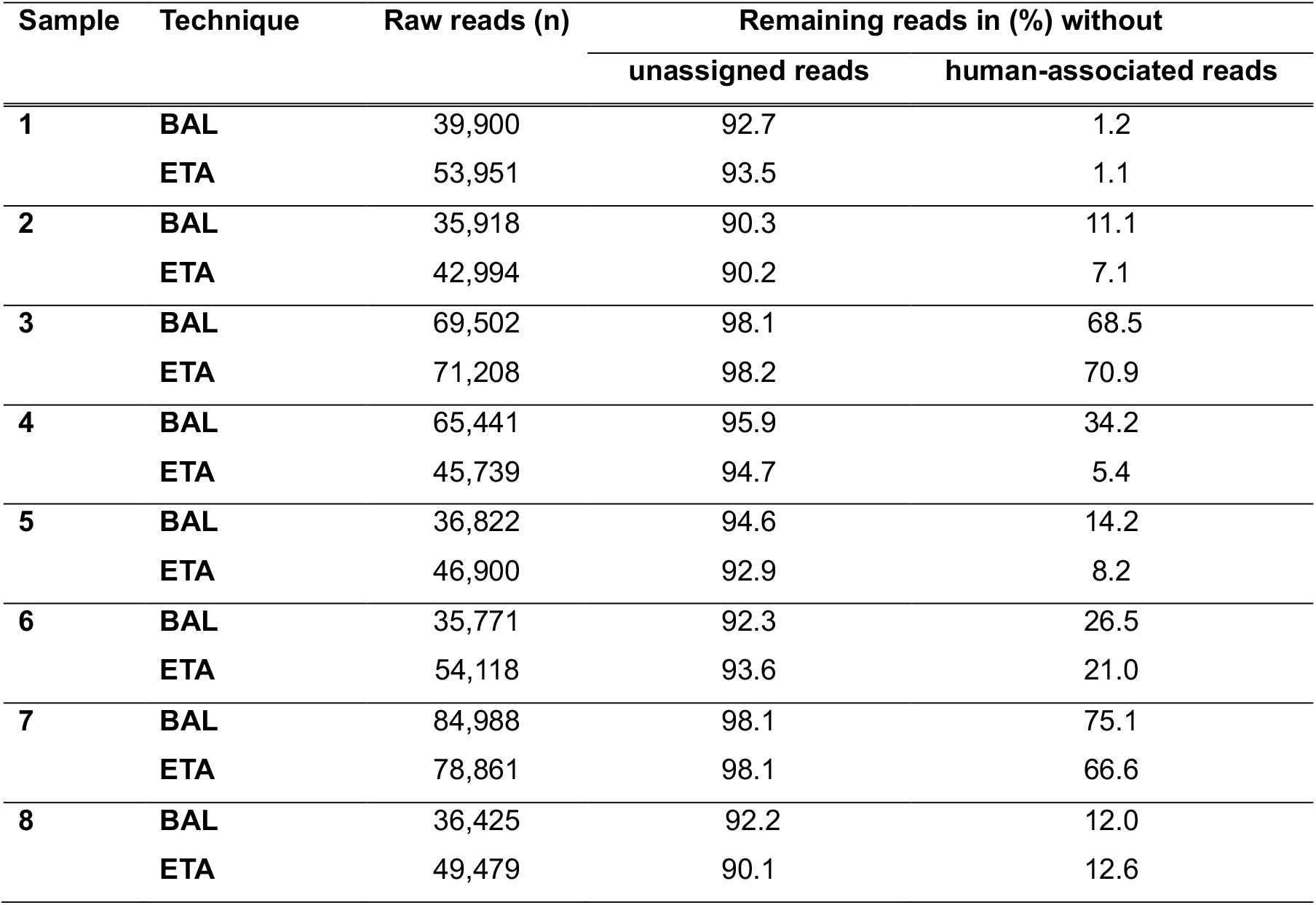
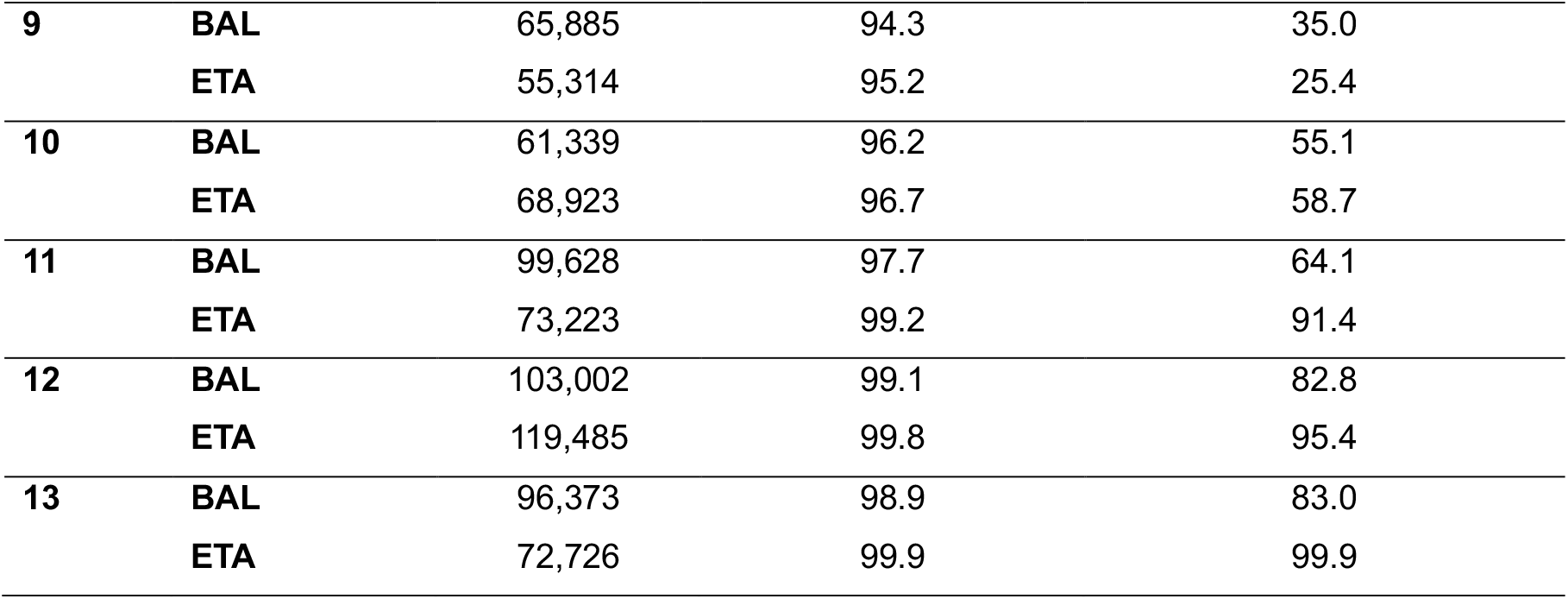
Number of raw reads and percentage of filtered reads detected in pairwise BAL and ETA samples.

**Figure 2.**
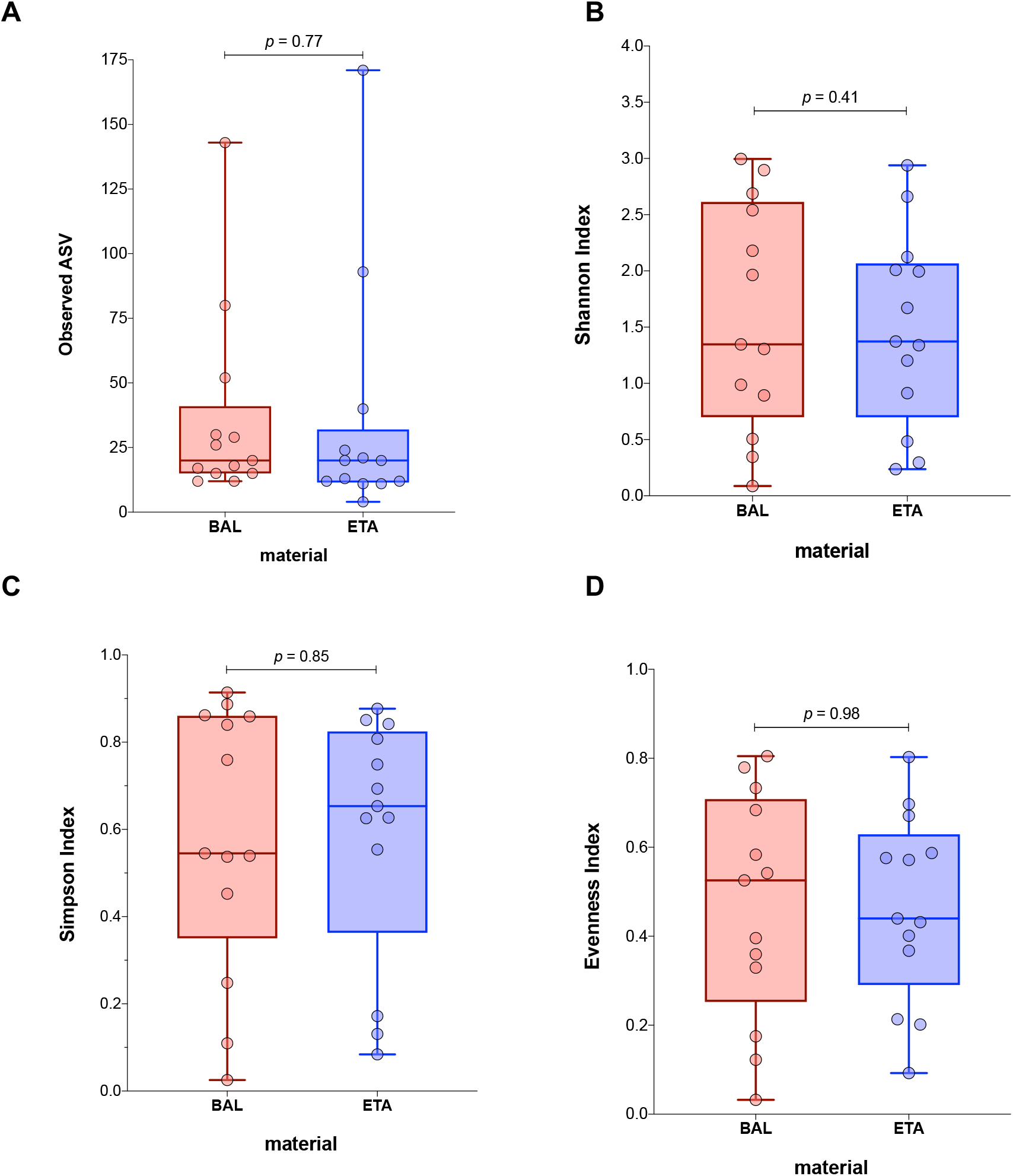
Comparison of species richness based on (A) observed ASV and evaluation of alpha diversity between BAL and ETA samples using (B) Shannon Index, (C) Simpson Index and (D) Evenness Index. Boxplots indicate minimum, 25^th^ percentile, median, 75^th^ percentile and maximum. Each data point represents an individual subject. Statistical analysis was performed using generalized linear mixed-effects models.

In addition, beta diversity based on Bray Curtis distance (*p* = 0.31) and Jaccard distance (*p* = 0.31) did not differ between BAL and ETA-derived samples (data not shown).

Since the objective of this study centered on the comparison of bacterial community structures between ETA and BAL specimens, the relative abundance of the 20 predominant taxa in each pairwise ETA and BAL samples was analyzed following adjustment for human sequences. The results of this systematic comparison are presented in **Figure 3**.

**Figure 3.**
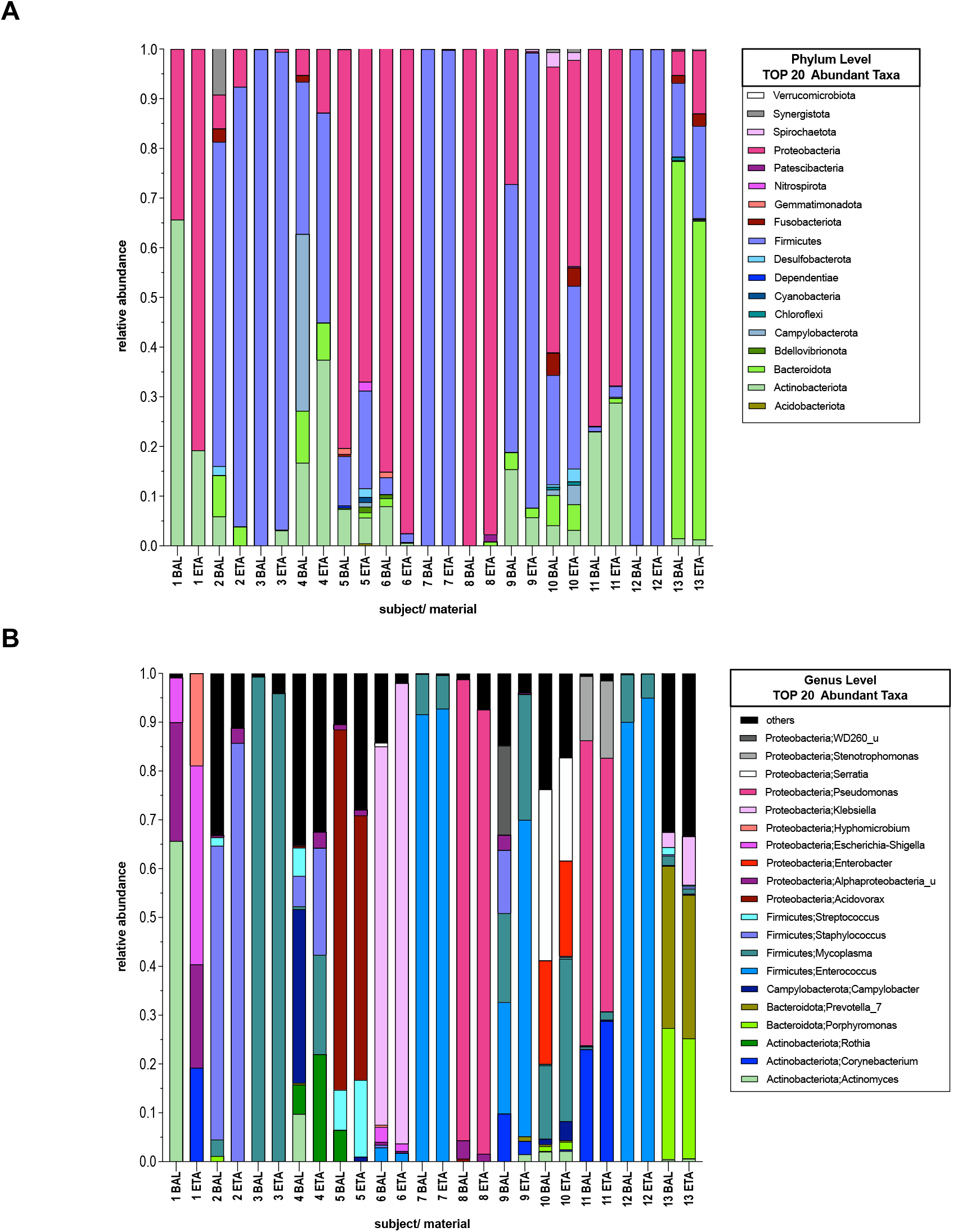
Fraction of relative abundance of the 20 major microbial taxa detected in pairwise BAL and ETA samples on. **(A). phylum level** **(B). genus level**. Legend colors correspondent to the most abundant microbiota identified in all subjects.

Bacterial community composition of the lung microbiome at the phylum level could be assigned to five major phyla which ranked *Firmicutes > Proteobacteria > Actinobacteriota > Bacteroidota > Campylobacterota* in each BAL and ETA (**Figure 3A**). Overall, a high resemblance between BAL and ETA was detected. Representative for the similarity of both specimens is the bacterial composition of *subject 3, 7* and *12* which was shaped predominantly by Firmicutes with a relative abundance of 99.9%, 99.9% and 99.8% in BAL as well as 96.2%, 99.8% and 99.9% in ETA, respectively. As second most common phylum Proteobacteria constituted 85.1%, 99.9% and 75.9% of BAL as well as 97.5%, 97.7% and 67.8% of ETA-derived bacterial community of *subject 6, 8* and *11*, respectively.

The seven most abundant taxa on genus level could be arranged by frequency as follows: *Enterococcus > Mycoplasma > others > Pseudomonas > Staphylococcus > Pseudomonas > Klebsiella* (**Figure 3B**).

We observed highly individual pattern among the 13 subjects, with increased overrepresentation of a single genus in the majority of critically-ill patients. As a case in point, the microbial composition of *subject 7* and *12* was enriched by *Enterococcus* comprising 91.6% and 90.0% of BAL, as well as 92.8% and 95.0% of ETA, respectively. Regarding further genera correspondent to *Firmicutes*, about 99.3% of taxa in BAL and 95.9% in ETA were assigned to *Mycoplasma* in *subject 3*. Furthermore, *Staphylococcus* (belonging to the phylum *Firmicutes)* accounted for 60.2% of reads in BAL and 85.8% in ETA in *subject 2*. With respect to *Proteobacteria*, high abundance of *Pseudomonas* was discovered in *subject 8* and *11* accounting for 94.4% and 62.5% of taxa in BAL, as well as 91.0% and 51.9% in ETA, respectively. In addition, *Klebsiella* was the most frequently detected genus in another pairwise set of specimens (*subject 6*) representing 77.5% of taxa identified in BAL and 94.3% of taxa collected from ETA.

Bacterial species that harbor the oral cavity as oral commensals, including members of *Actinomyces, Corynebacterium, Prevotella* and *Porphyromonas* were also detected occasionally in our cohort.

Unbalanced distribution of bacterial taxa was identified exclusively in *subject 1, 4* and *9*. Except for these three subjects, both sampling techniques exhibited similar abundance of microbiota.

To further elucidate the microbial composition of the lung microbiome, species richness was assessed based on ASV observed in BAL and ETA specimen (**Figure 4A**). The amount of detected ASV was 20.0 (*IQR* 15.0–41.0; *range*[*min*–*max*] 12.0–143.0) in BAL and 20.0 (*IQR* 11.5–32.0; *range*[*min*–*max*] 4.0–171.0) in ETA. Overall, the number of ASV identified in BAL and ETA did not differ significantly (*p* = 0.77) between both sampling methods.

**Figure 4.**
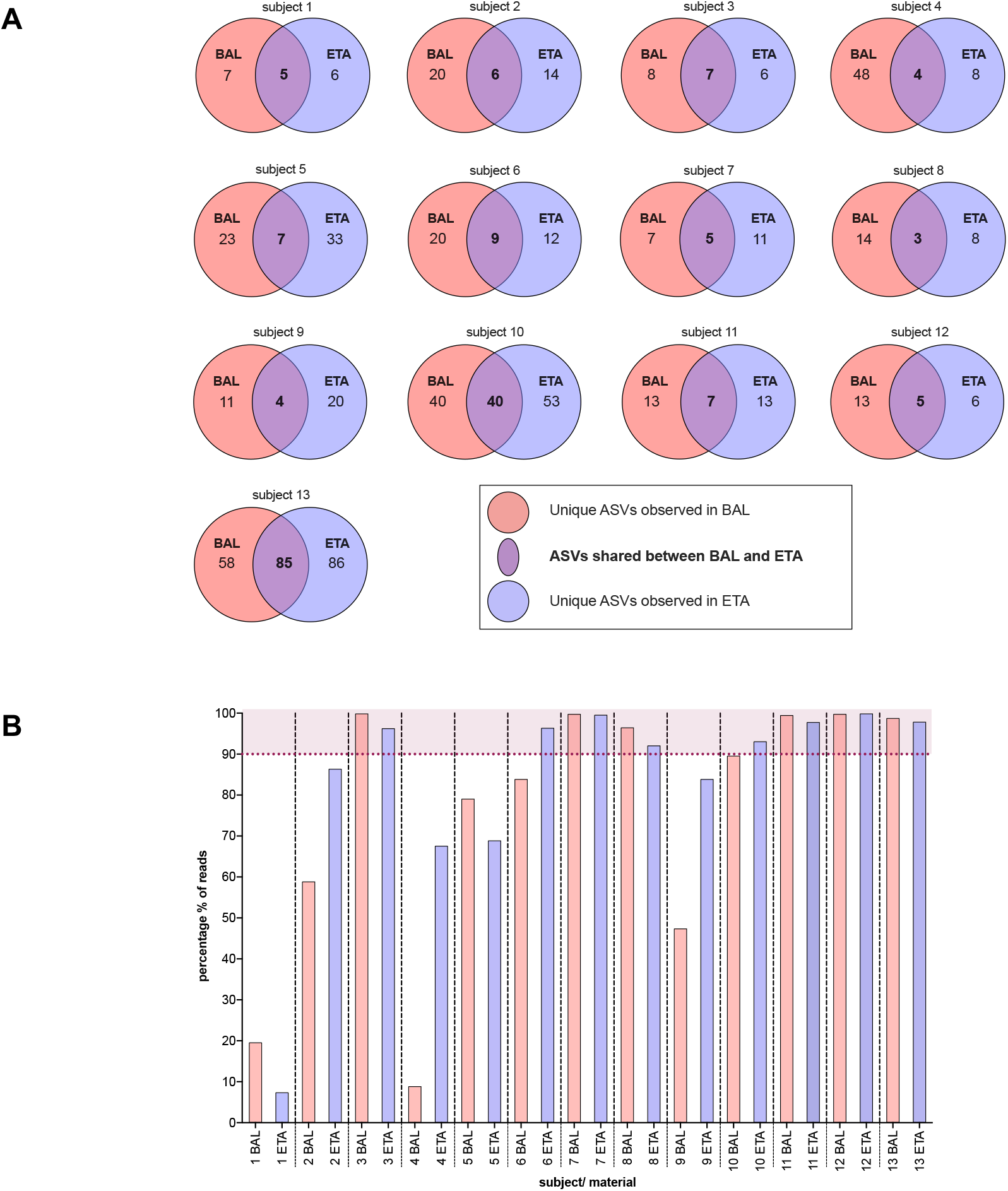
(A) Comparison of overlapping ASV between BAL and ETA per subject and (B) Percentage of reads per sample represented by these shared ASV. (A) Venn diagram of subjects 1-13 displays amount of unique ASV observed in BAL (red) and ETA (blue) as well as mutal ASV identified in both BAL and ETA (violet). According to their arrangement (subject 1-13), relative percentage of mutal ASV in BAL accounted for 41.7%, 23.1 %, 46.7%, 8.3%, 23.3%, 31.0%, 41.7%, 17.6%, 26.7%, 50,0%, 35,0%, 27.8% and 59.4% of total ASV. Relative percentage of shared ASV in ETA represented 45.5%, 30.0%, 53.8%, 33.3%, 17.5%, 42.9%, 31.3%, 27.3%, 16.7%, 43.0%, 35.0%, 45.5% and 49.7% of total ASV. (B) Percentage of reads constituting shared ASV in BAL (red) and ETA (blue) samples. Percentages >90.0% are marked violet.

**Figure 4A** shows the number of shared and unique ASV between both specimens per subject. Mutal ASV constituted 31.0% (*IQR* 23.2–44.2%; *range*[*min*–*max*] 8.3–59.4%) of observed ASV in BAL and 35.0% (*IQR* 28.7–45.5%; *range*[*min*–*max*] 16.7–53.8%) of total ASV in ETA.

Although the number of shared ASV for both specimens is mostly lower than that of the unique ASV, they account for a high percentage of reads in the respective samples with 89.6% (*IQR* 53.2–99.7%; *range*[*min*–*max*] 8.9–99.9%) in BAL and 93.1% (*IQR* 76.4– 97.9%; *range*[*min*–*max*] 7.4–99.9%) in ETA, as shown in **Figure 4B**.

In 46.2% (*n*= 6) of subjects the representation of ASV even exceeded 90% of reads. Solely in subject 1 and 4 the percentage of reads in BAL and ETA presented by ASV was scarce, with 19.6% and 7.4%, as well as 8.9% and 67.6%, respectively. As assumed, ASV in these subjects were mainly impacted by host contamination.

Overall, the percentage of detected ASV did not differ between both sampling techniques (**Figure 2A**, *p* = 0.77).

## DISCUSSION

Due to the lack of consensus on the microbial detection rate of ETA, we conducted an evaluation of identified respiratory microorganisms from pairwise ETA and BAL specimens of 13 critically ill patients based on 16S rRNA gene sequencing. In our study, the side-by-side evaluation of BAL and ETA samples indicated a profound resemblance at higher taxonomic levels; however, this methodological concordance declined progressively with increasing taxonomic resolution.

Specifically, alpha diversity supported by species richness (*p* = 0.77), Shannon diversity (*p* = 0.41), Simpson index (*p* = 0.85) and evenness (*p* = 0.98) did not differ between BAL and ETA. Furthermore, beta diversity based on Bray Curtis distance (*p* = 0.31) and Jaccard distance (*p* = 0.31) revealed comparable microbial community structures across both sampling techniques. In conclusion, the within-subject similarity at phylum and genus levels in our study supports that both sampling methods can equally reflect the lung microbiome at higher taxonomic ranks, complementing previous evidence in favor of their equivalence for microbial assessment [23–25].

Consistent with prior studies, the tracheobronchial microbiome predominantly consisted of the genus *Firmicutes, Proteobacteria* and *Bacteroidetes*, accompanied by minor proportions of *Actinobacteria* [5,26–30].

Apart from *subject 1, 4* and *9*, a strong taxonomic correlation was evident between BAL and ETA at phylum and genus levels. As pointed out previously, *subject 1* resulted in a low amount of remaining reads (*reads* < 1000) following filtering of human-associated sequences. Thus, the exclusion of subject 1 should be discussed. In *subject 4*, filtered reads and bacterial taxonomy were highly unbalanced between BAL and ETA. Despite a comparable number of raw reads, only 5.4% of reads remained for ETA in *subject 4* after quality filtering. Enhanced growth of *Rothia*, a bacterium commonly colonizing the oropharynx, was observed in ETA, whereas the detection of *Campylobacter* was confined to BAL [31,32]. Since isolation of Campylobacter in lower respiratory tract is rare, potential contamination during passage through the oropharynx should be acknowledged [33]. In *subject 9*, ETA was dominated by *Firmicutes* (90.5% of taxa), in particular *Enterococcus* and *Mycoplasma*, while BAL showed greater diversity, with *Firmicutes* accounting for 54% of taxa and the presence of *Proteobacteria*. This may reflect topographic variations between upper and lower airway segments. However, the presence of the oral commensal *Corynebacterium* (belonging to *Actinobacteriota*) in BAL could also indicate bronchoscope carryover or transient aspiration from the oral cavity [7,34,35].

Overall, ventilated patients exhibited considerable inter-individual heterogeneity in taxonomic distribution at both the phylum and genus levels [36,37]. Pathogens constituted the majority of detected pulmonary microbiota in both sample types, indicating lung dysbiosis in this critically ill cohort [7,26,36,38]. Patient-specific pathogens, such as *Mycoplasma, Klebsiella* and *Pseudomonas*, were equally overrepresented in ETA and BAL, likely reflecting underlying pulmonary infections [7,26,27]. Additionally, the overabundant growth of the genus *Pseudomonas* may also relate to biofilm formation on airway tubes with subsequent colonization of the associated airway [39]. Due to missing clinical data, the interactions between pathognomonic microbiota, subsequent respiratory disease and its diagnostic value were not assessed.

As expected, genera of the oral cavity, such as *Prevotella*, were also detected occasionally among our specimens [4,26,32,37,40]. Since the pulmonal microbiome is continuously shaped by microaspiration, these genera may be part of the common lung microbiome as well [6,7]. Historically, ETA-derived cultures have been deemed less reliable due to potential contamination with microbiota from the upper airway and oral cavity [9,41–44]. In our cohort, identification of oral commensals was confined to sporadic instances with ETA and BAL demonstrating similar abundance of oral commensals at higher taxonomic ranks, thus, challenging studies that advocate for the clear advantage of BAL [13,14,41–43].

To this date, the diagnostic accuracy of ETA compared to BAL has been targeted predominantly by conventional culture-based analyses, almost exclusively addressing cases of ventilator-associated pneumonia (VAP). In previous prospective studies comparative results and similar antibiotic resistance patterns were evident in both ETA and BAL-derived cultures collected from patients with VAP [24,45–47]. In addition, a systematic review reported no clinical advantage of invasive diagnostics with subsequent quantitative cultures over noninvasive diagnostics with nonquantitative cultures in suspected VAP [48]. In contrast, a major meta-analysis of pneumonia diagnosis acknowledged ETA-derived culture to be associated with higher sensitivity and low specificity compared to BAL [49]. A prospective study also preferred BAL over ETA in diagnosis of VAP due to ETA’s inferior positive predictive value, risking inappropriate antibiotic treatment and resistance [50]. Similarly, the current European Respiratory Society (ERS) guidelines recognized culture collected from invasive BAL as gold standard for the precise identification of causative organisms in patients with VAP [44].

However, conventional culture techniques are inherently restricted to cultivable microorganisms capable of growth under specific incubation conditions and medium, and thus may fail to accurately reflect the broad spectrum of the respiratory microbiome due to selectively favored bacterial growth [1,52]. Culture-independent methods, such as 16S rRNA gene sequencing, have broaden the detection range of organisms in the respiratory setting [1,3,18,34,35,52]. While culture-based methods remain the gold standard for initial pathogen identification [53], the adjunctive value of molecular diagnostics should be acknowledged, particularly in complex or culture-negative cases, polymicrobial infections or when detecting rare taxa in low-biomass samples that may be missed by conventional culture methods.

A prospective Taiwanese cohort study which compared microbiota profiles of BAL and ETA from patients with severe pneumonia observed a high correlation in bacterial community composition between the two specimens based on 16S rRNA amplicon sequencing method [29]. Consistent with Cheng et al. [29], our findings indicated similar microbial compositions between BAL and ETA at higher taxonomic levels, with no index differing significantly between both sampling methods (p > 0.05). Elaborating this aspect even further, Cheng et al. characterized ETA as a favorable procedure for identification of pathogens in pneumonia with less invasiveness and high accuracy [29].

Likewise, the establishment of shotgun metagenomic next-generation sequencing (mNGS) as a promising diagnostic technique has further challenged the lack of confidence in the diagnostic value of ETA. In contrast to conventional cultures that are limited to bacteria and fungi, mNGS facilitates the rapid and sensitive detection of a wider range of microorganisms including bacteria, viruses and fungi simultaneously [15,54]. A retrospective study reported positive rates of ETA and BAL pathogen detection of 96.7% and 80.7%, respectively, in patients with severe pneumonia, [54]. Notably, a prospective cohort study by Kalantar et al. emphasized the potential of ETA to even exceed BAL if molecular diagnostic methods are applied [55]. Contrarily, another retrospective study favored BAL mNGS due to significantly improved rates of confirmed pathogens and pneumonia [15].

Although 16S rRNA gene sequencing outperforms conventional culture methods in detecting a broader range of microbial taxa, it exhibits lower taxonomic resolution, particularly among closely related species [56–58].Therefore, full length 16S rRNA gene sequencing or metagenomic approaches are favored for accurate species-level identification [58,59], though not yet routinely implemented in clinical practice. As anticipated, the method comparability between BAL and ETA declined with enhanced taxonomic resolution (**Figure 4A**). Even though this observed ASV variability may be attributable solely to natural fluctuations of very rare or low-abundance taxa in low-biomass samples and microbial profiles of BAL and ETA samples may progressively align over time, the absence of longitudinal data precludes any conclusion on this matter. Moreover, the identification of recurrent ASVs and their assignment to phylogenetic trees was not feasible in this cross-sectional study, thereby limiting species-level resolution.

Due to the limited knowledge of the real in vivo diversity of the lower respiratory tract microbiome, it remains uncertain whether BAL or ETA yields a more precise representation at finer taxonomic resolutions. Therefore, further research is essential to fully elucidate this issue. In this context, our study may provide a methodological basis for future investigations addressing the lung microbiome.

With regard to the persistent controversy surrounding the detection rate of ETA, our results provide evidence supporting its equivalence to BAL for sequence-based microbial profiling at the phylum and genus levels. Thus, this research may facilitate rational guidance towards less invasive ETA as first-choice procedure for lower respiratory tract monitoring in routine clinical practice at an intensive care unit, when genus-level classification is appropriate.

Our research is subject to certain limitations. As a monocentric study, the sample size is limited and recruitment of healthy controls was not feasible due to ethical considerations. Besides, the characterization of the lung microbiome was restricted solely to bacterial taxa. The anonymous character of this study prevented acquisition of clinical and personal data, such as physical constitution, medical history and administration of drugs, so potential confounders remain, though we mitigated this by paired sampling of BAL and ETA from the same patient and focusing on pairwise analysis.

No concurrent sampling of crucial contamination sources (e.g. endotracheal tube, bronchoscope rinse, skin, laboratory reagents) as controls was performed and thus, potential contamination cannot be ruled out.

Contrary to prior research centered on VAP patients, our study comprised a broader population of critically ill patients, enhancing generalizability. Finally, the effects of antibiotic or antimycotic treatment, systemic steroids and immunosuppressive therapy on the lung microbiome were not assessed, though its administration and adjustments may have further altered the composition of the respiratory microbiome.

## CONCLUSION

Our study indicates that ETA provides equivalent taxonomic information to BAL at higher taxonomic levels in an unselected cohort of mechanically ventilated patients. These results confirm the suitability of ETA as a reliable alternative to BAL for microbial profiling in routine clinical practice, particularly when genus-level resolution is adequate. However, considering the anonymous design of this study, further research is required to assess the potential impact of sampling techniques on clinical outcomes and the dynamics of respiratory microbiota during antibiotic and antimycotic treatment.

## Abbreviations

ASV: amplicon sequence variant
BAL: bronchoalveolar lavage
DNA: Desoxyribonucleic Acid
e.g.: exempli gratia/for example
ETA: endotracheal aspirate
ERS: European Respiratory Society
Fr: French
IQR: Interquartile Range
mL: microliter
mNGS: metagenomic next-generation sequencing
nM: nanometer
PCR: polymerase chain reaction
r: ribosomal
RNA: Ribonucleic Acid
sec: seconds
VAP: ventilator-associated pneumonia
µl: microliter

## Author Contributions

Conceptualization, S.P., U.L..; Sample collection, J.K., T.K., N.L., U.L.; Data curation, S.P., S.G., U.L.; Formal analysis, S.P., S.G.; Methodology, S.P., S.G., U.L.; Resources, S.G., U.L.; Supervision, S.G., U.L.; Visualization, S.P., S.G.; Writing—original draft, S.P., S.G., U.L.; Writing—review and editing, S.G., U.L., M.S., N.L., J.K., T.K.

**All authors have read and agreed to the published version of the manuscript**.

## Institutional Review Board Statement

The local ethic committee confirmed no objections concerning publication.

## Data Availability Statement

The data presented in this study are available on request from the corresponding author.

